# High-Potency Polypeptide-based Inhibition of Enveloped-virus Glycoproteins

**DOI:** 10.1101/2021.04.05.438537

**Authors:** Jianpeng Ma, Adam Campos Acevedo, Qinghua Wang

## Abstract

Specific manipulation of proteins post-translationally remains difficult. Here we report results of a general approach that uses a partial sequence of a protein to efficiently modulate the expression level of the native protein. When applied to coronavirus, human immunodeficiency virus, Ebolavirus, respiratory syncytial virus and influenza virus, polypeptides containing highly conserved regions of the viral glycoproteins potently diminished expression of the respective native proteins. In the cases of coronavirus and influenza virus where multiple strains were tested, the polypeptides were equally effective against glycoproteins of other coronavirus and influenza strains with sequence identity as low as 27%, underscoring their high insensitivity to mutations. Thus, this method provides a platform for developing high-efficacy broad-spectrum anti-viral inhibitors, as well as a new way to alter expression of essentially any systems post-translationally.

## Introduction

Proteins constitute the building blocks of the cells. Changes in native proteins including mutations, altered functions or levels of expression are fundamental to many forms of diseases including cancer and neurodegenerative diseases, just to name a few. Therefore, manipulation of proteins in a highly specific manner at the post-translational level in the cells is presumably the most straightforward approach for treating diseases. However, this remains a difficult task.

To address these unresolved challenges, here we report a new approach in which a partial sequence of a native protein can efficiently modulate its expression level. Application of this method to the highly conserved regions of glycoproteins from coronavirus, human immunodeficiency virus, Ebolavirus, respiratory syncytial virus and influenza virus resulted in potent polypeptide-based inhibition. Strikingly, the polypeptides developed for SARS-CoV-2 coronavirus and influenza A/H3N2 virus were equally effective against glycoproteins of other coronavirus and influenza strains with sequence identity as low as 27%, underscoring their high insensitivity to mutations. Considering that infectious diseases persist to be a major medical burden around the globe, as exemplified by the ongoing coronavirus disease 2019 (COVID-19) pandemic caused by severe acute respiratory syndrome coronavirus 2 (SARS-CoV-2), this method provides a platform for developing high-efficacy broad-spectrum anti-viral inhibitors. Furthermore, this approach represents a new way to alter expression of essentially any systems post-translationally.

## Results

### The concept of polypeptide-based interference using coronavirus as an example

Viral membrane fusion proteins such as coronavirus spike proteins are oligomeric Class-I transmembrane glycoproteins on the viral envelope^1^. Coronavirus spike (S) proteins are cleaved to give rise to N-terminal S1 regions and C-terminal S2 regions (**Fig.1a**). The N-terminal S1 regions are the major target for neutralizing antibodies elicited by natural infection or vaccination, and therefore under constant positive selection for escape variants. For instance, since the onset of COVID-19, four major variants (α-δ) of SARS-CoV-2 with extensive mutations have emerged. The mutations on SARS-CoV-2 spike (SARS2-S) proteins^2-4^ were concentrated in the S1 regions **(Fig.1a)**, leading to tighter binding with host cell surface receptor, human angiotensin-converting enzyme 2 (hACE2), higher virulence, resistance to antibody neutralization, and partial escape from natural infection or vaccine induced sera^5-9^. On the other hand, the C-terminal S2 regions responsible for oligomerization and membrane fusion^10-14^ are more conserved among different coronavirus strains. Upon entry into a host cell, the viral genome will guide the synthesis of new coronavirus spike proteins by ribosomes, which are then folded, assembled and translocated into endoplasmic reticulum (ER) membranes and transit through the ER-to-Golgi intermediate compartment for interaction with newly replicated genomic RNA to produce new virions^15,16^.

**Figure 1.**
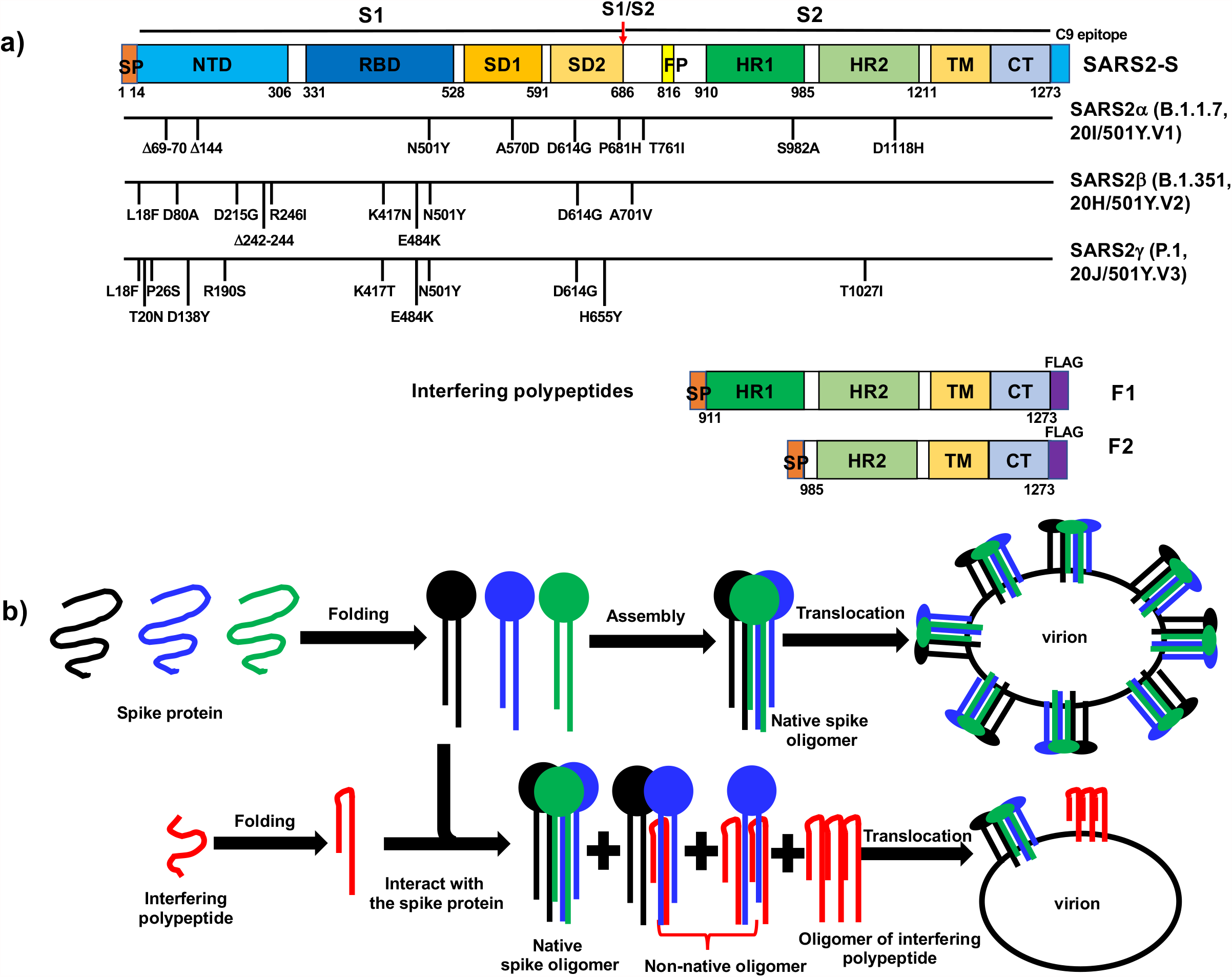
The concept of polypeptide-based protein interference against coronavirus spike proteins. **a)**. Domain organization of COVID-19 SARS2-S, the mutations in recent SARS2-S variants (SARS2α-S, SARS2β-S, and SARS2γ-S) and the design of interfering polypeptides F1 and F2. SP: Signal peptide; NTD: N-terminal domain; RBD: receptor-binding domain; SD1: subdomain 1; SD2: subdomain 2; FP: fusion peptide; HR1: heptad repeat 1; HR2: heptad repeat 2; TM: transmembrane domain; CT: Cytoplasmic tail. The cleavage at S1/S2 (red arrow) gives rise to N-terminal S1 fragment and C-terminal S2 fragment. The signal peptide sequence at the extreme N-termini of F1 and F2 allowed the polypeptides to be translocated in the same way as COVID-19 SARS2-S. At the extreme C-termini, SARS2-S had a C9 epitope recognized by C9-rhodopsin antibody 1D4, while both F1 and F2 had a FLAG-tag. **b)**. Diagram of polypeptide-based interference targeting coronavirus spike proteins. Top row: in the normal situation, the spike proteins are synthesized, folded and formed native spike oligomers, which are anchored on virion envelope. Bottom row, interfering polypeptides form non-native oligomers with the wild-type spike proteins, thus reducing the level of native spike oligomers on the envelope of new virions.

We hypothesized that stably foldable fragments of the coronavirus spike proteins, for instance, polypeptides derived from the SARS2-S S2 regions, which maintain the same oligomeric interface as the native spike proteins, would form non-native oligomers with the wild-type spike proteins, thus significantly lowering the level of native spike oligomers on the envelope of new virions, and impairing their infectivity (**Fig.1b**). More importantly, the fragment derived from the conserved S2 region would be more resistant to mutations and could potentially be a pan-coronavirus inhibitor. To test this hypothesis, we made two polypeptides derived from the SARS2-S S2 sequence, F1 and F2 (**Fig.1a, Fig.S1, S2**). F1 polypeptide encompassed amino-acid residues 911∼1273, while F2 harbored residues 985∼1273. Both polypeptides contained a N-terminal signal peptide (SP) as the wild-type SARS2-S for cell surface translocation.

### F1 polypeptide potently inhibited the expression and surface translocation of multiple coronavirus spike glycoproteins

We first assessed the impacts of F1 or F2 on the expression and cell surface translocation of SARS2-S protein. Transient transfection of SARS2-S-coding plasmid resulted in a good level of proteins detected in HEK293T whole cell lysate, with most of the expressed proteins were cleaved (**Fig.2a**). This high-efficiency cleavage of SARS2-S proteins agreed with the novel polybasic cleavage site at the S1/S2 boundary^10,11,17,18^. Moreover, only the S2 fragments of the cleaved SARS2-S protein were labeled by biotin, affinity-purified by anti-biotin antibody and detected in the cell surface fraction (**Fig.2a**), suggesting that only properly cleaved SARS2-S protein were translocated to cell surface. Most impressively, when F1-coding plasmid was co-transfected with SARS2-S-coding plasmid, even at a twofold molar ratio, the predominant cleaved S2 band of SARS2-S was almost completely diminished in whole cell lysate and in cell surface fraction (**Fig.2a**). Thus, F1 strongly inhibited the expression and surface translocation of SARS2-S. In sharp contrast, F2 did not exhibit any significant interference even at a tenfold molar ratio (**Fig.2a**).

**Figure 2.**
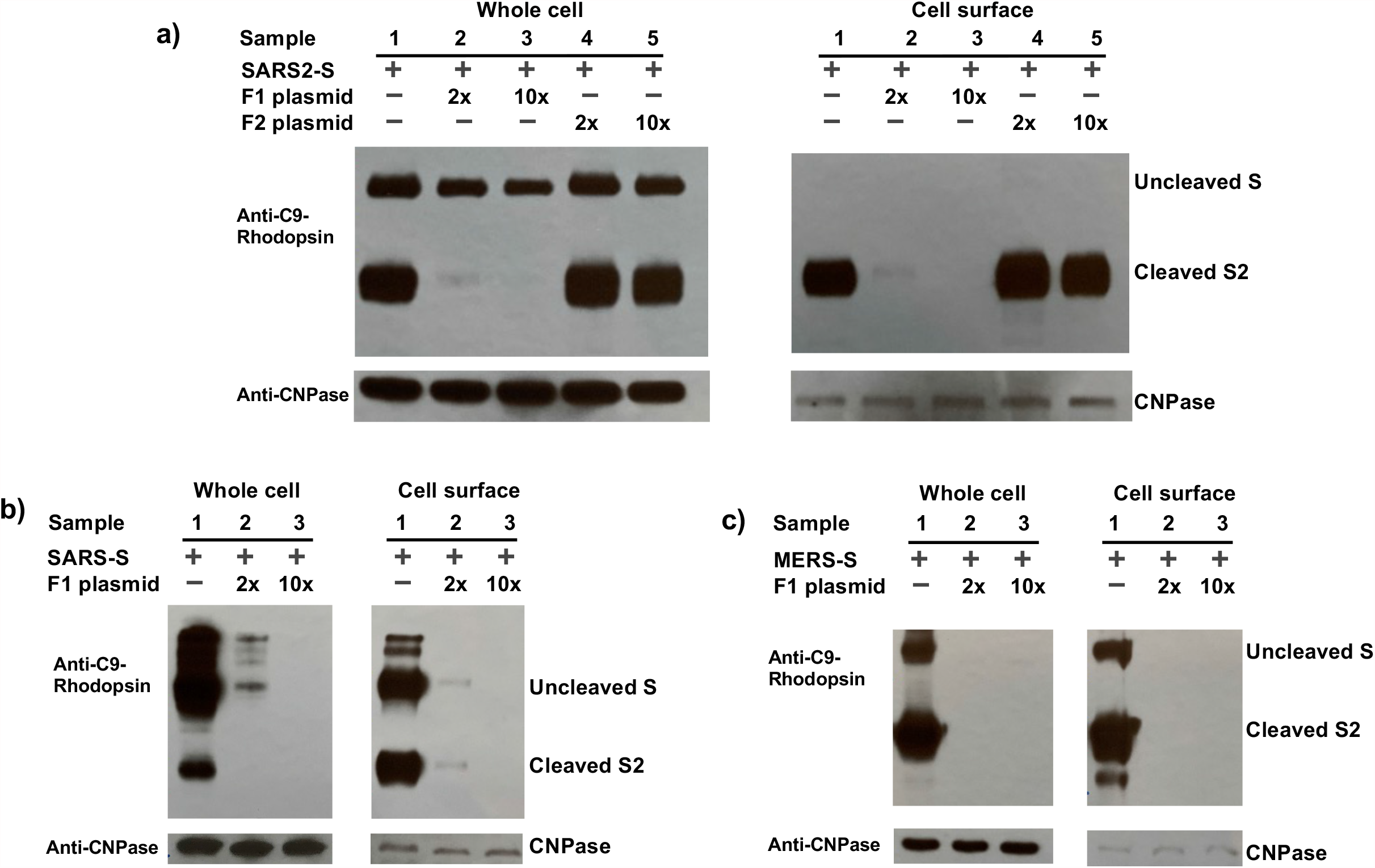
F1 significantly reduced expression and surface translocation of three coronavirus spike glycoproteins. **a)**. COVID-19 SARS2-S. **b)**. 2002 SARS-S. **c)**. 2012 MERS-S. For panels **a-c)**, the levels of S protein in whole cell lysate (left) or in cell surface fraction (right) were compared for HEK293T cells transfected with S-coding plasmid only (Sample 1), or together with twofold (Sample 2) or tenfold molar ratio (Sample 3) of F1-coding plasmid, and in panel **a)** with twofold (Sample 4) or tenfold molar ratio (Sample 5) of F2-coding plasmid. Endogenous membrane-anchored protein CNPase detected by anti-CNPase antibody was used as an internal control.

Besides COVID-19 SARS-CoV-2, the past 20 years have witnessed two other major threats from coronaviruses, the 2002-2003 outbreak from severe acute respiratory syndrome coronavirus (SARS-CoV), and the 2012-2014 outbreak from Middle East respiratory syndrome coronavirus (MERS-CoV)^19,20^. Sequence comparison of 2002 SARS-CoV spike (SARS-S), 2012 MERS-CoV spike (MERS-S) and COVID-19 SARS2-S uncovered a wide range of sequence identity levels among them (**Fig.S2, S3**). For instance, compared with full-length COVID-19 SARS2-S, 2002 SARS-S was 77% amino-acid sequence identical, while 2012 MERS-S was only 35% (**Fig.S2, S3**). If only considering the regions included in F1 polypeptide, the identical residues became 94% for 2002 SARS-S and 42% for 2012 MERS-S (**Fig.S2, S3**). These spike proteins were thus ideal for testing the sensitivity of F1-induced inhibition to amino-acid mutations. Strikingly, even with F1-coding plasmid at a twofold molar ratio, the full-length SARS-S and MERS-S bands and the cleaved S2 bands were almost completely diminished in whole cell lysate and in cell surface fraction (**Fig.2b**,**c**). Consistent with the robust inhibitory activity exhibited by F1 (**Fig.2**), a high level of F1 polypeptide was constantly detected in cell surface fraction (**Fig.S4a-c**). In marked contrast, a limited level of F2 polypeptide was detected in cell surface fraction (**Fig.S4a**), in agreement with the non-interference of F2 (**Fig.2a**).

### F1 interfered with coronavirus spike proteins at the protein level

In order to probe the mechanism of F1-mediated inhibition, we analyzed the mRNA levels of coronavirus spike proteins in the absence or presence of F1 or F2-coding plasmids (**Fig.S5, S6**). Clearly co-transfection of F1- or F2-coding plasmids did not significantly change the mRNA levels of coronavirus spike proteins, which were kept at a relatively constant level (between 50%∼150%) comparing to the samples that were transfected only with plasmids coding the respective coronavirus spike proteins (**Fig.S5**). In sharp contrast, the mRNA levels of F1 were at the order of 2^6.5^∼2^12^ when normalized against endogenous GAPDH (**Fig.S6**), justifying the high potency of F1-mediated interference. The mRNA levels of F2 were comparable with those of F1 (**Fig.S6a**), in marked contrast with the low level of F2 polypeptide detected in cell surface fraction (**Fig.S4a**), suggesting that the non-inhibition of F2 (**Fig.2a**) was likely due to the instability of the synthesized F2 polypeptide. These data suggested that F1-mediated inhibition of various coronavirus spike glycoproteins was not at the mRNA level.

We then investigated whether F1-mediated inhibition is at the protein level *via* direct interaction with SARS2-S to form non-native oligomers in the cell. SARS2-S and F1 were each tagged with a monomeric green fluorescent protein (GFP) variant at the extreme C-terminus, CFP for SARS2-S (termed as SARS2C) and YFP for F1 (named as F1Y) (**Fig.S7**). Since the expression of SARS2-S was completely diminished when F1- and SARS2-S-coding plasmids were co-transfected at a 2:1 molar ratio (**Fig.2a**), we used F1Y- and SARS2C-coding plasmids at a reduced 1:1 molar ratio to transiently transfect HEK293T cells, and monitored fluorescence resonance energy transfer (FRET) ratio between them by using the three-cube approach^21^. Although the non-native oligomers formed by SRARS2C and F1Y were expected to be highly unstable, FRET signals between them were robustly detected (**Fig.S7**), supporting direct formation of non-native oligomers between SARS2-S and F1.

### F1 minicircle potently interfered with the expression and surface translocation of all three coronavirus spike glycoproteins

The high potency of F1 in inhibiting the expression and surface translocation of spike glycoproteins from human coronaviruses that caused severe outbreaks or pandemic between 2002 to 2021 suggests that F1 has a high promise to become an effective therapeutic agent against different coronavirus lineages over a long time period. Therefore, we sought to identify a convenient way to deliver F1 for therapeutic purpose. Since F1 directly interacts with its target spike protein to form non-native oligomers concomitant with protein synthesis and folding, F1-coding gene needs to be delivered to the site of action. Minicircles are a type of newly developed DNA carriers for gene therapy^22^. The main features of minicircles include the cleaner gene background with minimal viral or bacterial gene elements, sustained high-level protein expression, and more importantly, the small size that may allow the use of aerosols for drug delivery^23^. The latter may be a distinct advantage against coronavirus-caused respiratory diseases.

We made a F1 minicircle by inserting the F1-coding sequence into the parental minicircle cloning vector pMC.CMV-MCS-SV40polyA (**Fig.3a**), and tested its efficacy in inhibiting the expression and surface translocation of coronavirus spike glycoproteins. Compared to the controls where no minicircle was used, the presence of F1 minicircles, at a merely 4.5-fold molar ratio, almost completely abolished cell surface translocation of all three spike proteins (**Fig.3b-d**). It is important to note that this level of inhibition was achieved under the situation where pcDNA3.1-based plasmids harboring coronavirus spike-coding genes were efficiently replicated in HEK293T cells, while F1 minicircle cannot.

**Figure 3.**
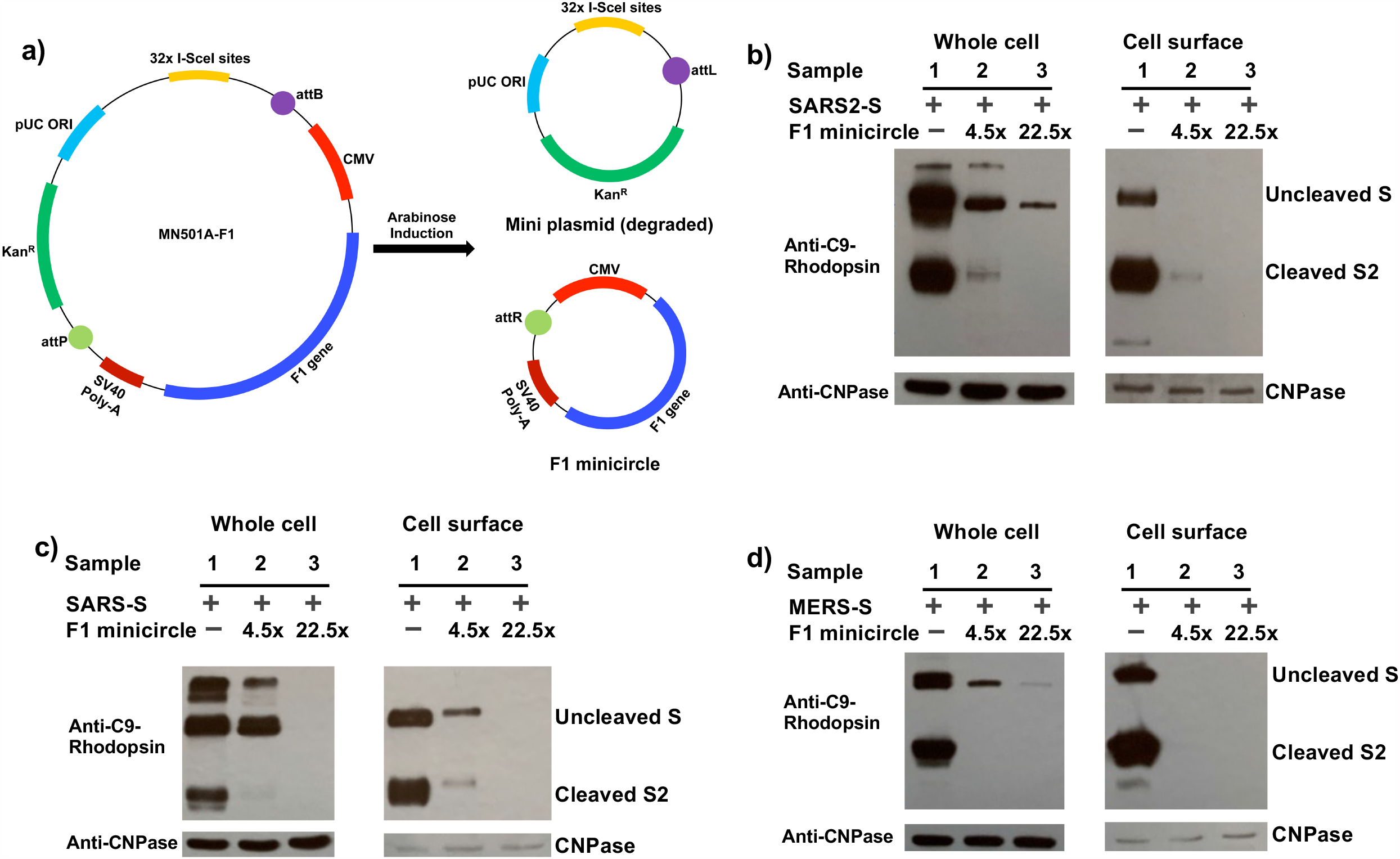
F1 minicircle significantly reduced expression and surface translocation of three coronavirus spike glycoproteins. **a)**. Diagram for the production of F1 minicircle used in this study. **b)**. COVID-19 SARS2-S; **c)**. 2002 SARS-S; **d)**. 2012 MERS-S. The levels of uncleaved S protein and the cleaved S2 protein in whole cell lysate (left) or in cell surface fraction (right) were compared for HEK293T cells transfected with S-coding plasmid only (Sample 1), or together with 4.5-fold (Sample 2) or 22.5-fold molar ratio (Sample 3) of F1 minicircles. Endogenous membrane-anchored protein CNPase detected by anti-CNPase antibody was used as an internal control.

### F1 minicircle reduced the level of SARS2-S protein on intact pseudoviruses and impaired pseudovirus infectivity

To investigate the consequences of the reduced cell surface translocation of coronavirus spike proteins by F1 minicircle, we compared the level of SARS2-S protein on pseudoviruses generated using luciferase-expressing, *env*-defective HIV-1 genome plasmid pRL4.3-Luc-R^-^E^-^ in the presence of different molar ratios of control minicircle made from the empty parental vector (termed as MN501A) or F1 minicircle. In order to make sure that only spike proteins anchored on the pseudovirus envelope were accounted for, we employed QuickTiter Lentivirus Titer kit to precipitate intact pseudoviruses from cleared supernatant prior to analysis by western blot. Impressively, even with a twofold molar ratio of F1 minicircle, almost no SARS2-S was detected on the generated intact pseudoviruses (**Fig.4a**). Not surprisingly, these pseudoviruses completely failed to infect hACE2-expressing HEK293T cells (**Fig.4b**).

**Figure 4.**
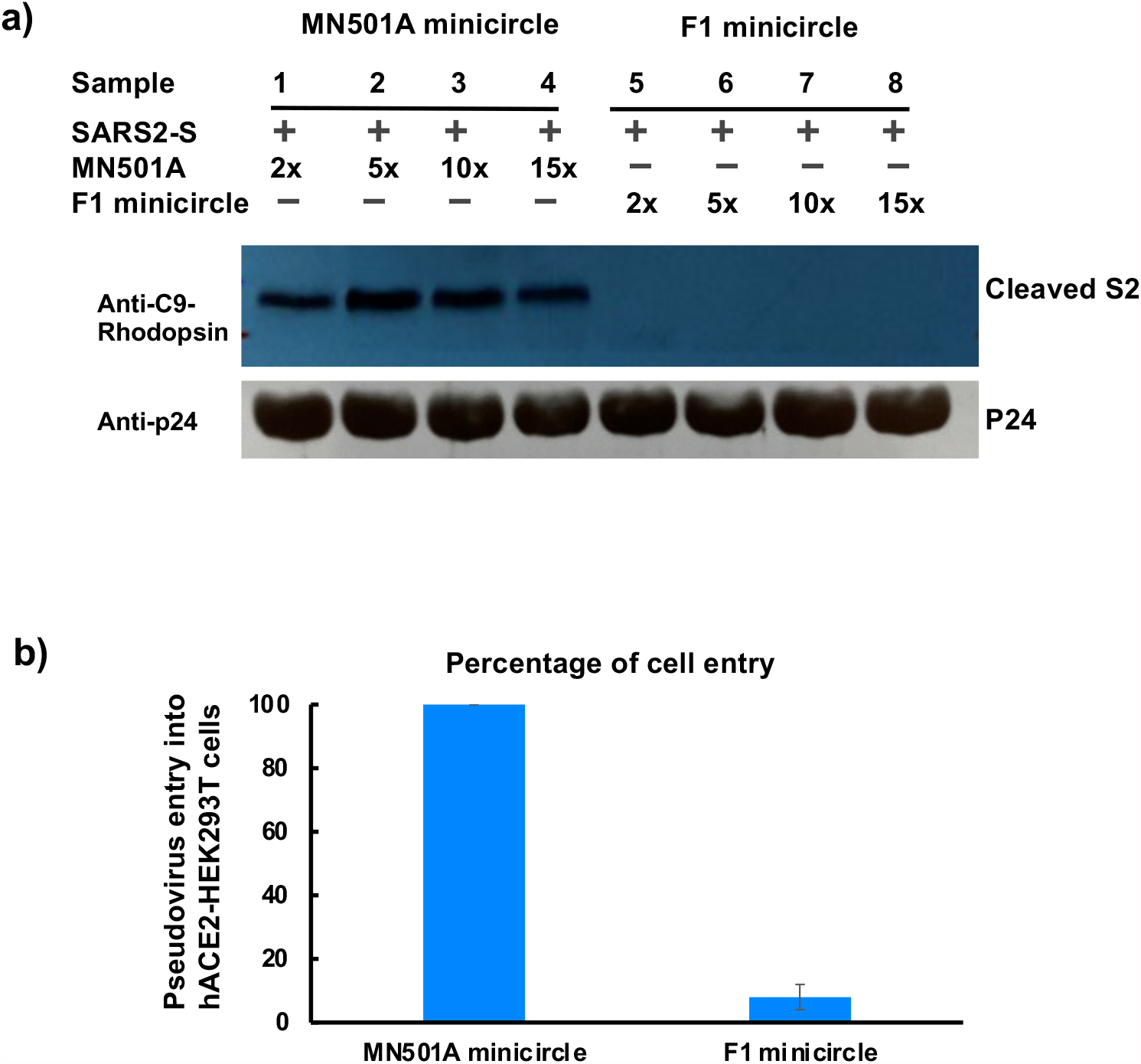
F1 minicircle significantly reduced the level of SARS2-S on intact pseudoviruses and impaired pseudovirus infectivity. **a)**. The level of cleaved S2 protein was compared for intact pseudovirus produced from HEK293T cells transfected with SARS2-S-coding plasmid and different ratios of empty minicircle control, MN501A (Sample 1∼4), or F1 minicircle (Sample 5∼8). p24 detected by anti-p24 antibody was used as an internal control. **b)**. Pseudovirus generated in the presence of F1 minicircle at a twofold molar ratio completely lost the ability to infect hACE2-expressing HEK293T cells. The infectivity of pseudovirus generated in the presence of MN501A minicircle at a twofold molar ratio was considered as 100%.

### Targeted inhibition of glycoproteins from other enveloped viruses

Using the same strategy, we designed polypeptide-based inhibitors for human immunodeficiency virus-1 (HIV-1) envelope glycoprotein (gp160), Zaire ebolavirus (EBOV) glycoprotein (GP), human respiratory syncytial virus (RSV) A fusion protein (F) and influenza virus type A and B hemagglutinin (HA) (termed as gp160*i*, GP*i*, F*i* and HA*i*, respectively, where the letter “*i*” denotes their inhibitory activities, **Fig.S1**). When the inhibitor-coding plasmids were co-transfected with the corresponding glycoprotein-coding plasmid, at 5∼15-fold molar ratio, the corresponding glycoprotein was almost completely diminished in whole cell lysate and in cell surface fraction (**Fig.5a-d**). In particular, the HA*i* polypeptide derived from the HA sequence of the 2019-2020 flu vaccine strain A/Hong Kong/45/2019 (H3N2) virus was able to potently inhibit the expression and translocation of HA proteins of all four flu vaccine strains, A/Hawaii/70/2019 (H1N1), A/Hong Kong/45/2019 (H3N2), B/Washington/02/2019 (B/Victoria lineage (B/Vic)), B/Phuket/3073/2013 (B/Yamagata lineage (B/YM)) (**Fig.5d**). It is worth noting that A/Hong Kong/45/2019 (H3N2) HA shares a sequence identity of 43%, 27% and 29% with the HA protein from A/Hawaii/70/2019 (H1N1), B/Washington/02/2019 (B/Vic) and B/Phuket/3073/2013 (B/YM), respectively (**Fig.S3**). If only considering the interfering polypeptide HA*i* region, the sequence identity was at 53%, 33% and 34%, respectively. These data further demonstrated that the use of a partial native sequence is an effective method for targeted reduction of protein expression of these important human pathogens, even when the level of sequence identity is as low as 27%.

**Figure 5.**
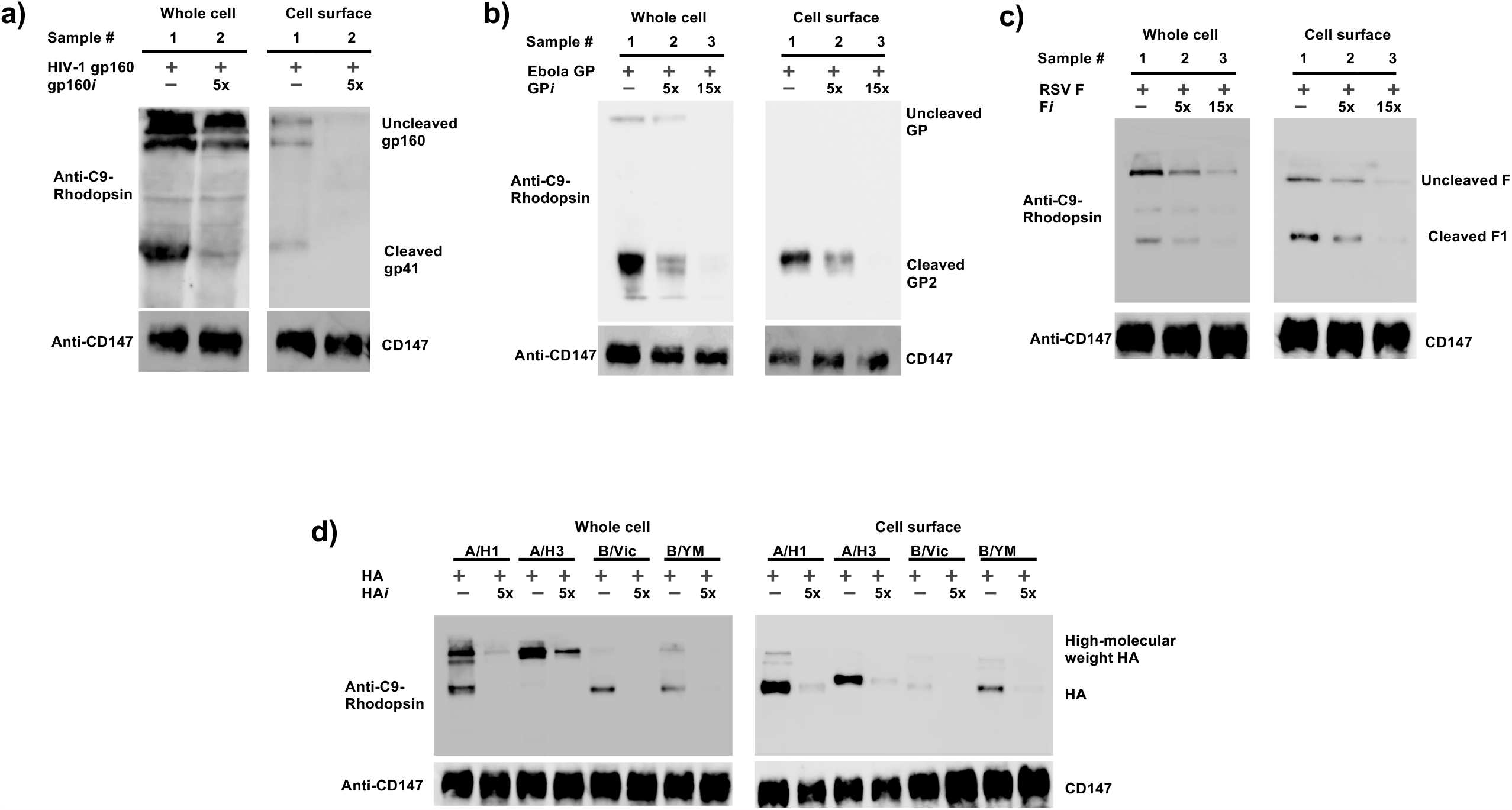
Inhibitory polypeptides for other enveloped-virus glycoproteins. **a)**. HIV-1 gp160. **b)**. EBOV GP. **c)**. RSV F. **d)**. Influenza HAs. For panels **a-d**), the levels of envelope glycoproteins in whole-cell lysate (left) or in cell-surface fraction (right) were compared for HEK293T cells transfected with envelope-coding plasmid only, or together with 5∼15-fold molar ratio inhibitor-coding plasmid. Endogenous CD147 protein detected by anti-CD147 antibody was used as an internal control.

## Discussion

In the current research, we present results of a new approach that uses a partial sequence of a protein to efficiently modulate its expression level. In the case of coronaviruses, the F1 polypeptide derived from COVID-19 SARS2-S potently reduced the expression and surface translocation of the spike proteins from all three human coronaviruses that caused major regional outbreaks or global pandemic in the last 20 years, despite as low as 35% amino-acid sequence identity among them. Although extensive mutations were found on SARS2-S proteins in recent SARS-CoV-2 variants^2-4^ (**Fig.1a**), the regions corresponding to F1 polypeptide were highly conserved, as exemplified by the nearly identical sequence of SARS2α-S, a variant of SARS2-S, with F1 (**Fig.S3b**). Therefore, F1 polypeptide should be effective on the spike proteins of almost any emerging COVID-19 SARS-CoV-2 variants in the future. Furthermore, since the spike proteins of other human coronaviruses (including HCoV-HKU1 and HCoV-OC43, HCoV-NL63 and HCoV-229E) and 2012 MERS-CoV have a similar level of sequence identity with F1 (**Fig.S3b**), F1 polypeptide may be equally effective on all these spike proteins. Additionally, we demonstrated that the same strategy was also effective on glycoproteins from other enveloped viruses including HIV-1, EBOV, RSV and influenza viruses. Therefore, this polypeptide-based approach represents a new concept for targeted manipulation of viral and non-viral proteins with high potency and fine tunability.

## Materials and Methods

### Materials

The pcDNA3.1 plasmids harboring the genes coding SARS-CoV spike (GenBank accession number AFR58740.1), SARS-CoV-2 spike (GenBank accession number QHD43416.1), and human ACE2 (GenBank accession number NM_021804) were kind gifts from Dr. Fang Li^12^ (Addgene plasmid No. 145031, 145032 and 145033, respectively). The pcDNA3.1 plasmids harboring the genes coding MERS-CoV spike (GenBank accession number QBM11748.1), F1 or F2 polypeptides, pEZT-BM plasmids harboring the genes coding HIV-1 gp160 (GenBank accession number NP_057856.1); Zaire EBOV GP (GenBank accession number AAN37507.1); RSV F (GenBank accession number QKN22797.1); HA proteins for A/Hawaii/70/2019 (H1N1), A/Hong Kong/45/2019 (H3N2), B/Washington/02/2019 (B/Vic), and B/Phuket/3073/2013 (B/YM)), gp160*i*, GP*i*, F*i* and HA*i* were synthesized by Genscript Biotech (Piscataway, NJ, USA). The parental minicircle vector pMC.CMV-MCS-SV40polyA (Cat. No. MN501A-1), ZYCY10P3S2T *E. coli* minicircle producer strain^24^ competent cells (Cat. No. MN900A-1) and Arabinose Induction Solution (Cat. No. MN850A-1) were purchased from System Biosciences (Palo Alto, CA, USA). The luciferase-expressing, *env*-defective HIV-1 genome plasmid pRL4.3-Luc-R^-^E^-^ (Cat. No. 3418) was obtained through the NIH AIDS Reagent Program, Division of AIDS, NIAID, NIH, USA. C9-rhodopsin antibody 1D4 HRP (Cat. No. sc57432 HRP), 2’,3’-cyclinc nucleotide 3’-phosphodiesterase (CNPase) antibody (Cat. No. A01308), and CD147 antibody (Cat. No. ab108308) were purchased from Santa Cruz Biotechnology, Inc. (Dallas, TX, USA), Genscript Biotech (Piscataway, NJ, USA), and Abcam (Cambridge, UK), respectively. QuickTiter Lentivirus Titer kit (Cat. No. VPK-107) was purchased from Cell Biolabs Inc (San Diego, CA, USA). Pierce Cell surface Protein Biotinylation and Isolation Kit (Cat. No. A44390) was purchased from Thermo Scientific, Waltham, MA, USA. Quick-RNA miniprep Kit (Cat. No. R1054) and ZymoPURE II Plasmid Maxiprep Kit (Cat. No. D4203) were obtained from Zymo Research, Irvine, CA, USA. iScript Reverse Transcription Supermix (Cat. No. 1708840) was purchased from Bio-rad, Hercules, CA, USA. Bimake SYBR Green qPCR Master Mix (Cat. No. B21203) was obtained from Bimake, Houston, TX, USA. Lipofectine 3000 (Cat. No. L3000015) was obtained from Invitrogen, Carlsbad, CA, USA. One-Glo EX Luciferase Assay System (Cat. No. E8110) was purchased from Promega, Madison, WI, USA. HEK293T cells (Cat. No. CRL-11268) were purchased from American Type Culture Collection (Manassas, VA, USA) and maintained in DMEM medium containing 10% fetal bovine serum (FBS) at 37°C incubator supplied with 5% CO2.

### Cell surface biotinylation and protein purification

Cell surface biotinylation and protein purification were performed using Pierce Cell surface Protein Biotinylation and Isolation Kit following the manufacturer’s instruction. Briefly, cell surface proteins on HEK293T cells were first labeled with Sulfo-NHS-SS-Biotin at 4°C for 30 minutes, which were then stopped by adding Tris-buffered saline and further washed. After cells were lysed with Lysis Buffer, lysate was cleared by centrifugation. Cleared lysate was incubated with NeutrAvidin Agarose to allow binding of biotinylated proteins. After extensive wash, the bound proteins were eluted with Elution Buffer containing 10 mM DTT. The cleared lysate (“Whole cell” fraction) and eluted proteins (“Cell surface” fraction) were run on 10% SDS-PAGE and the spike proteins were detected by C9-rhodopsin antibody 1D4 HRP. Endogenous membrane-anchored protein CNPase detected by anti-CNPase antibody or CD147 detected by anti-CD147 antibody were used as an internal control.

### Total RNA isolation and RT-qPCR

Total RNA was purified using Quick-RNA miniprep Kit. Reverse transcription was carried out using iScript Reverse Transcription Supermix. qPCR was performed using Bimake SYBR Green qPCR Master Mix with the following primers:

1060 (SARS-S Forward): GTTCAAGGACGGCATCTACTT

1061 (SARS-S Reverse): ACGCTCTGGGACTTGTTATTC

1062 (SARS2-S Forward): GACAAAGTGCACCCTGAAGA

1063 (SARS2-S Reverse): GGGCACAGGTTGGTGATATT

1089 (MERS-S Forward): GAACGCCTCTCTGAACTCTTT

1090 (MERS-S Reverse): GTCCTCGGTGATGTTGTATGT

1091 (F1 and F2 Forward): GATTAGAGCCGCCGAGATTAG

1092 (F1 and F2 Reverse): GGACTGAGGGAAAGACATGAG

Since the synthesized genes of F1 and F2 were optimized for mammalian expression and different from the gene coding for SARS2-S, the qPCR primers 1091 and 1092 were unique to F1 and F2, while 1062 and 1063 were unique to SARS2-S.

### Minicircle production

F1-coding gene was cloned into the minicircle parental vector pMC.CMV-MCS-SV40polyA (MN501A) to yield MN501A-F1. MN501A or MN501A-F1 was transformed into ZYCY10P3S2T *E. coli* minicircle producer strain competent cells following the manufacturer’s instruction. The production of minicircle DNA was induced with the addition of Arabinose Induction Solution. Minicircle DNA was purified by using ZymoPURE II Plasmid Maxiprep Kit per the manufacturer’s instruction.

### Pseudovirus generation, precipitation and concentration

Pseudovirus generation followed the protocol reported earlier^25^. Essentially, HEK293T cells were seeded on 6-well plates the night before. The next day, pcDNA3.1-SARS2-S (0.6 μg) and pRL4.3-Luc-R^-^E^-^ (0.6 μg) were used to transfect one-well HEK293T cells using Lipofectine 3000. MN501A minicircle or F1 minicircle at indicated molar ratio was included in the transfection mixture. At 16 hours post-transfection, the HEK293T cells were fed with fresh medium. At 48 hours after medium change, the supernatant of each well of the 6-well plates was harvested, and centrifuged at 300 g for 5 minutes to remove cell debris. Intact pseudoviruses were purified using QuickTiter Lentivirus Titer kit following the manufacturer’s instruction. The virus lysate was analyzed by western blot using rhodopsin antibody 1D4 HRP for spike proteins, and FITC-conjugated anti-p24 mAb and HRP-conjugated anti-FITC mAb for p24, which served as an internal control. A portion of pseudovirus-containing supernatant was concentrated by PEG8000 and used for luciferase assay of cell entry.

### Luciferase assay of cell entry by pseudoviruses

HEK293T cells were seeded on 100 mm dishes the night before. The next day, HEK293T cells were transfected with 10 μg pcDNA3.1-hACE2 using Lipofectine 3000. At 16 hours post-transfection, the cells were resuspended in FBS-free DMEM medium, and plated onto 96-well white plates to which 10 μL concentrated pseudoviruses was already added to each well. Two hours later, each well was added with an equal volume of DMEM containing 20% FBS. The cells were further incubated for 36 hours, then an equal volume of One-Glo EX Luciferase Assay Reagent was added, after incubation for 3 minutes, the luminescence signals were recorded.

### FRET between SARS2C and F1Y

High quality/high resolution automated imaging was performed on a GE Healthcare DVLive epifluorescence image restoration microscope using an Olympus PlanApoN 60X/1.42 NA objective and a 1.9k x 1.9k pco.EDGE sCMOS_5.5 camera with a 1024×1024 FOV. The filter sets used were: CFP (438/24 excitation, 470/24 emission) and YFP (513/17 excitation, 559/38 emission). Donor and Acceptor control channels were acquired using CFP-CFP and YFP-YFP, respectively. FRET images were acquired using CFP-YFP excitation and emission filter pair. Cells were chosen with similar intensity profiles in donor and acceptor prior to acquisition while under 37°C and 5% CO_2_ environmental conditions. Z stacks (0.25µm) covering the whole cell (∼12µm) were acquired before applying a conservative restorative algorithm for quantitative image deconvolution using SoftWorx v7.0, and saving files as a max pixel intensity projection tiff for each individual channel. The FRET ratio was determined by using the three-cube approach^21^ that was defined as the ratio of YFP emission in the presence of FRET over that in the absence of FRET:

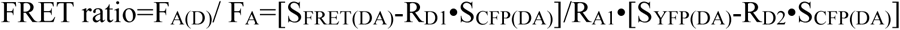

Where R_D1_= S_FRET(D)_/S_CFP(D)_; R_D2_=S_YFP(D)_/S_CFP(D)_; R_A1_=S_FRET(A)_/S_YFP(A)_; and S_FRET_, S_CFP_ and S_YFP_ with D, A, DA in parenthesis refer to fluorescence signals of the FRET, CFP or YFP channel when only SARS2C (containing CFP as donor (D)), F1Y (containing YFP as acceptor (A)), or both SARS2C and F1Y-coding plasmids (both donor and acceptor (DA)) were included in the transfection. Averages from untransfected cells were subtracted from same-day fluorescence values for each filter cube. According to the definition, FRET ratio=1.0 means no FRET, while FRET ratio>1.0 indicates the presence of FRET between the proteins carrying donor and acceptor proteins.

## Acknowledgements

J.M. acknowledges support from the Welch Foundation (Q-1512). Q.W. thanks support from the Welch Foundation (Q-1826). A.A.C was partially supported by a postdoctoral training fellowship from the Computational Cancer Biology Training Program of the Gulf Coast Consortia (CPRIT Grant No. RP170593). Imaging used the Integrated Microscopy Core at Baylor College of Medicine and the authors thank Hannah Johnson and Dr. Fabio Stossi at Integrated Microscopy Core at Baylor College of Medicine for their help with collecting the FRET data.

## Author contributions

J.M. and Q.W. conceptualized the project; J.M. and Q.W. developed the methodology; J.M., A.A.C. and Q.W. performed the investigations; J.M. and Q.W. wrote the original draft; J.M., A.A.C. and Q.W. reviewed and edited the paper.

## Supplementary Information

**Figure S1.**
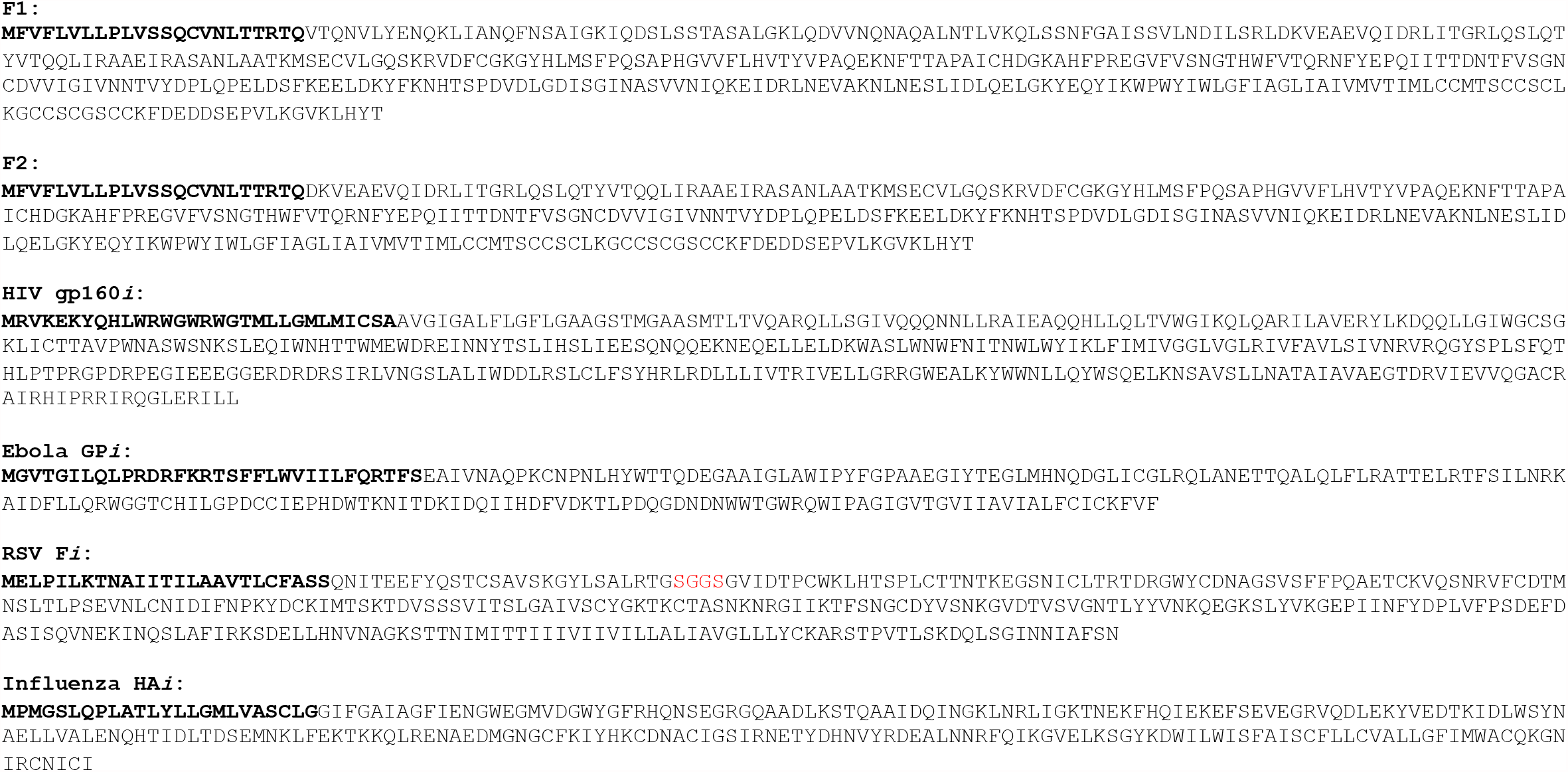
Amino-acid sequences of interfering polypeptides used in this study. Single-letter codes were used for amino acids. The signal peptide sequences were highlighted in boldface at the N-terminus. The sequences were derived from: SARS2-S (GenBank accession number AFR58740.1), HIV-1 gp160 (GenBank accession number NP_057856.1); Zaire EBOV GP (GenBank accession number AAN37507.1); RSV type A F (GenBank accession number QKN22797.1); and influenza virus A/HongKong/45/2019 (H3N2) HA. The residues SGGS in red color in RSV F*i* represent an introduced linker.

**Figure S2.**
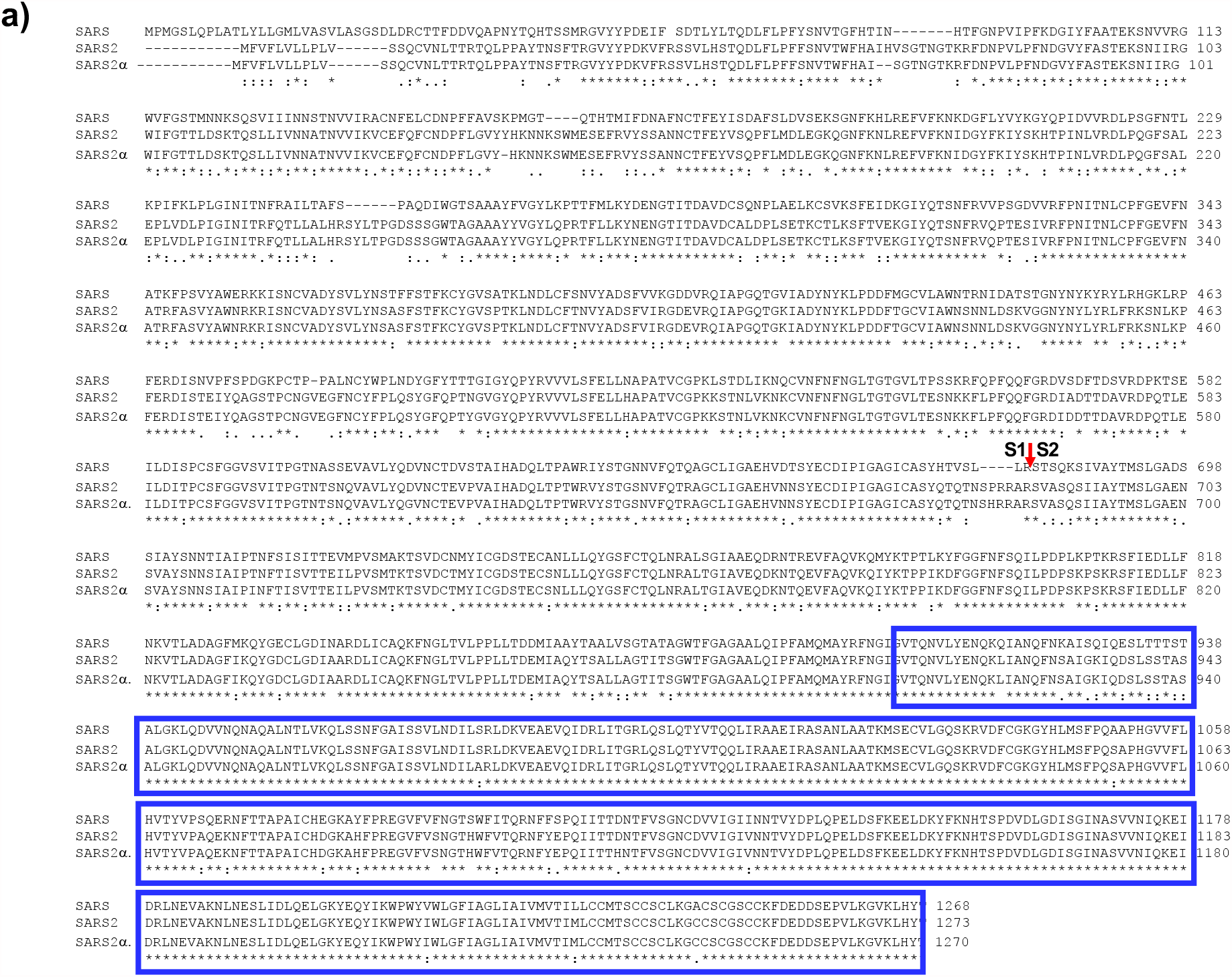
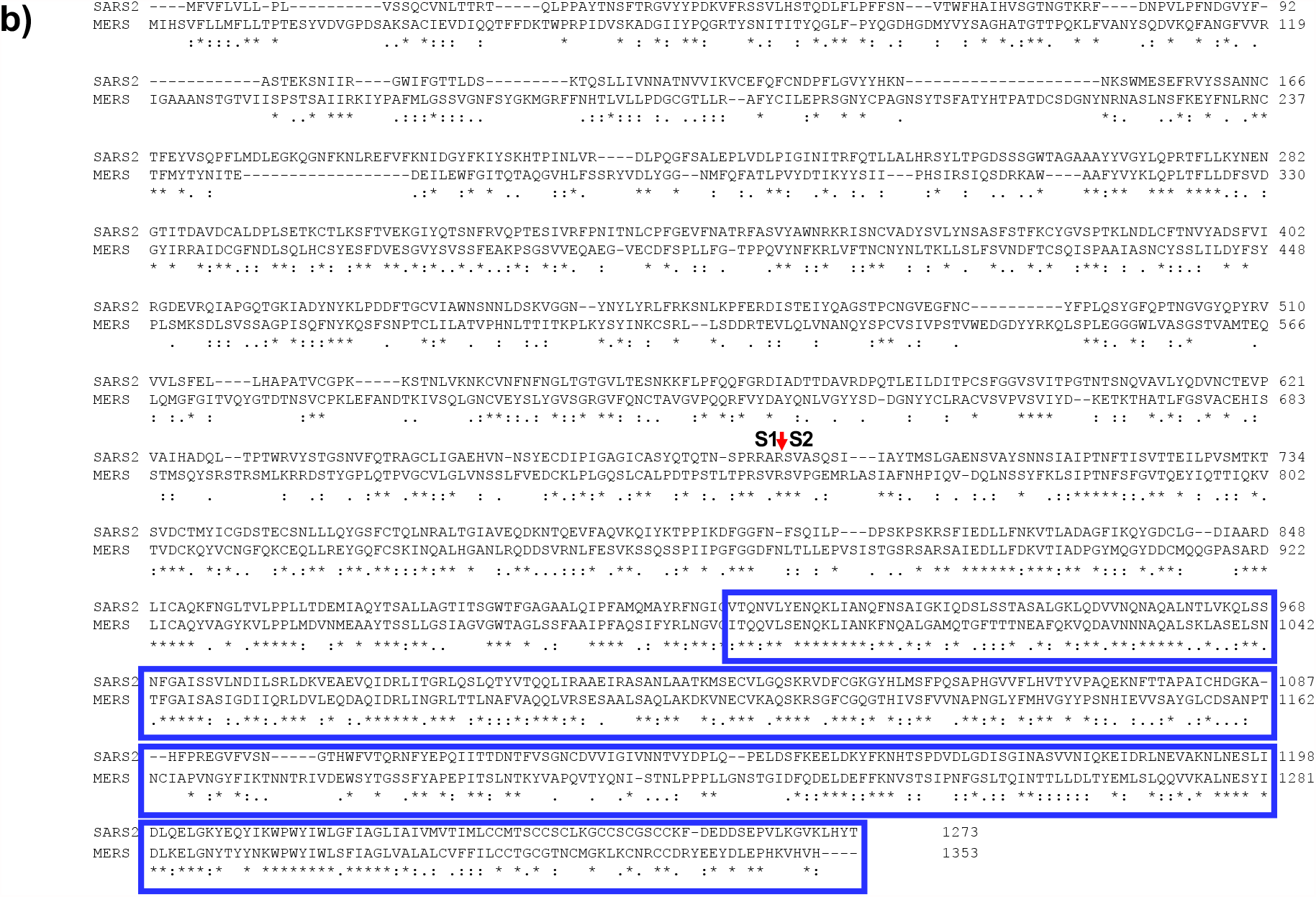
Sequence alignments of coronavirus spike proteins. **a)**. The alignment of SARS2-S (SARS2), SARS-S (SARS), and SARS2α-S (SARS2α). **b)**. The alignment of SARS2-S (SARS2) and MERS-S (MERS) proteins. In both panels, the range of residues included in the F1 polypeptide was highlighted in blue boxes. The cleavage site between S1 and S2 was shown by a red arrow. Multiple sequence alignments were performed using CLUSTAL Omega^24^. The sequences were: SARS2-S (GenBank accession number AFR58740.1), SARS-S (GenBank accession number QHD43416.1), SARS2α-S (GenBank accession number QQH18545.1), and MERS-S (GenBank accession number QBM11748.1).

**Figure S3.**
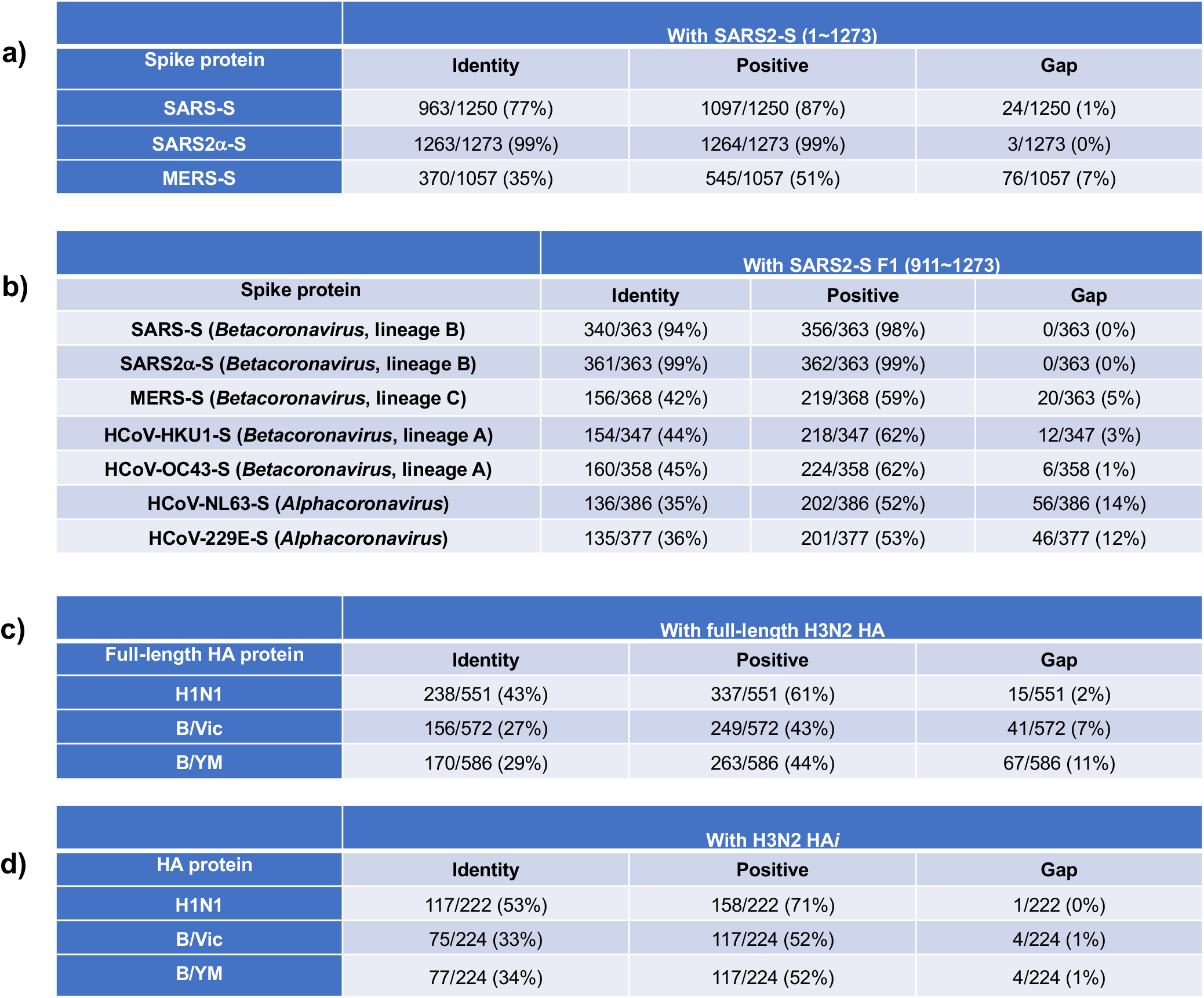
Sequence conservation of representative envelope glycoproteins. **a)**. The comparison of SARS-S, SARS2α-S and MERS-S proteins with full-length SARS2-S. **b)**. The comparison of representative human coronavirus spike proteins with SARS2-S F1 polypeptide (residues 911∼1273). **c)**. The comparison of H1N1, B/Vic and B/YM HA proteins with full-length H3N2 HA. **d)**. The comparison of H1N1, B/Vic and B/YM HA with H3N2 HA*i* polypeptide. In panels **a)** and **b)**, shown were the number of residues that were identical (Identity), similar (Positive) and unmatched (Gap) comparing to SARS2-S, the total residues compared (after “/”) and the corresponding percentage (in parenthesis). The total residues varied due to gaps during sequence alignments. The sequences were: SARS2-S (GenBank accession number AFR58740.1), SARS-S (GenBank accession number QHD43416.1), SARS2α-S (GenBank accession number QQH18545.1), MERS-S (GenBank accession number QBM11748.1), HCoV-HKU1-S (Uniprot accession number Q0ZME7), HCoV-OC43-S (Uniprot accession number P36334), HCoV-NL63-S (Uniprot accession number Q6Q1S2), and HCoV-229E-S (Uniprot accession number P15423). In panels **c)** and **d)**, the HA sequences were from the four flu vaccine strains in 2019-2020, A/Hawaii/70/2019 (H1N1), A/Hong Kong/45/2019 (H3N2), B/Washington/02/2019 (B/Vic), and B/Phuket/3073/2013 (B/YM).

**Figure S4.**
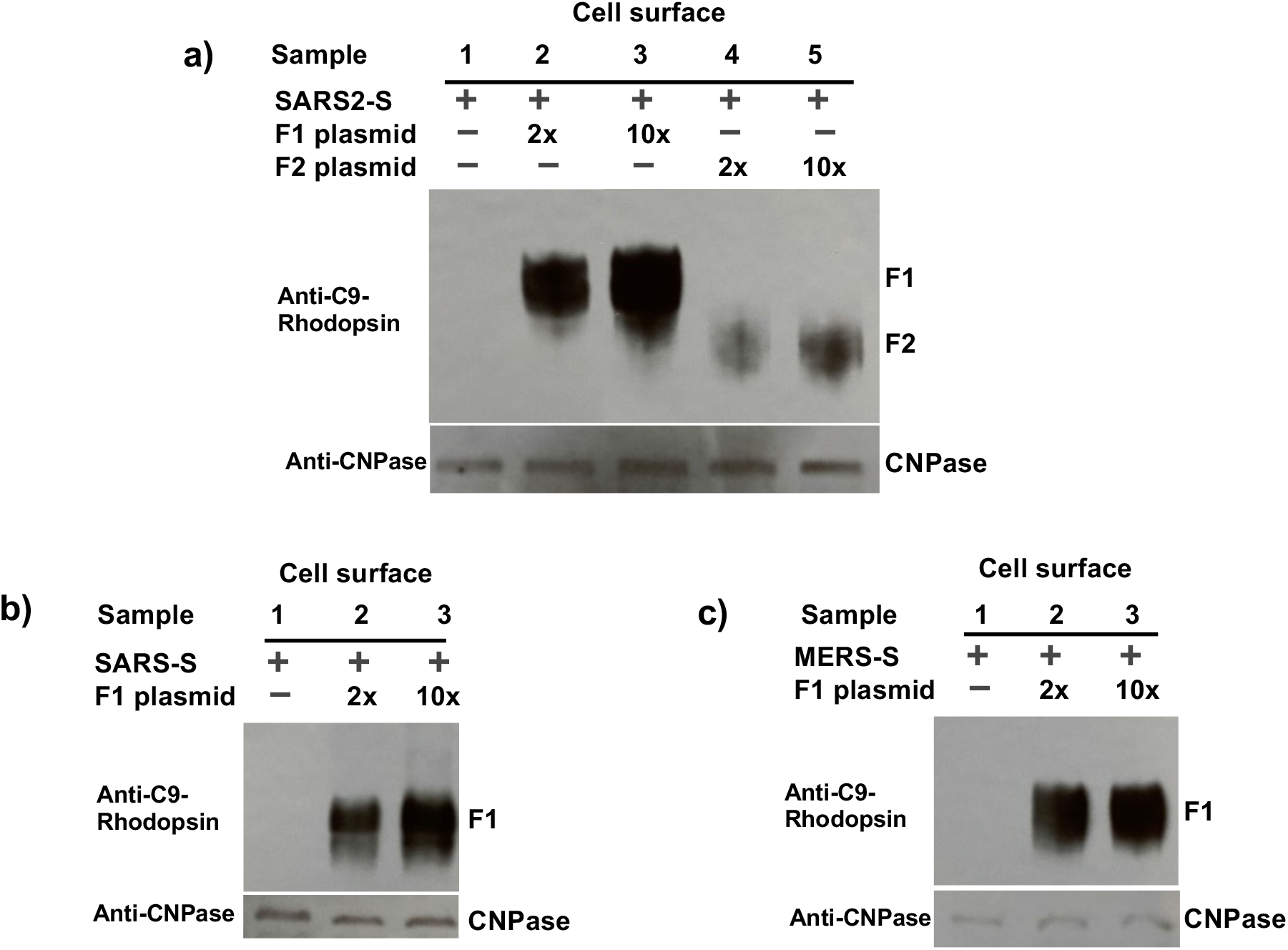
Protein expression levels of F1 and F2 when co-transfected with coronavirus spike protein-coding plasmids. **a)**. with COVID-19 SARS2-S; **b)**. with 2002 SARS-S; **c)**. with 2012 MERS-S. Endogenous membrane-anchored protein CNPase detected by anti-CNPase antibody was used as an internal control.

**Figure S5.**
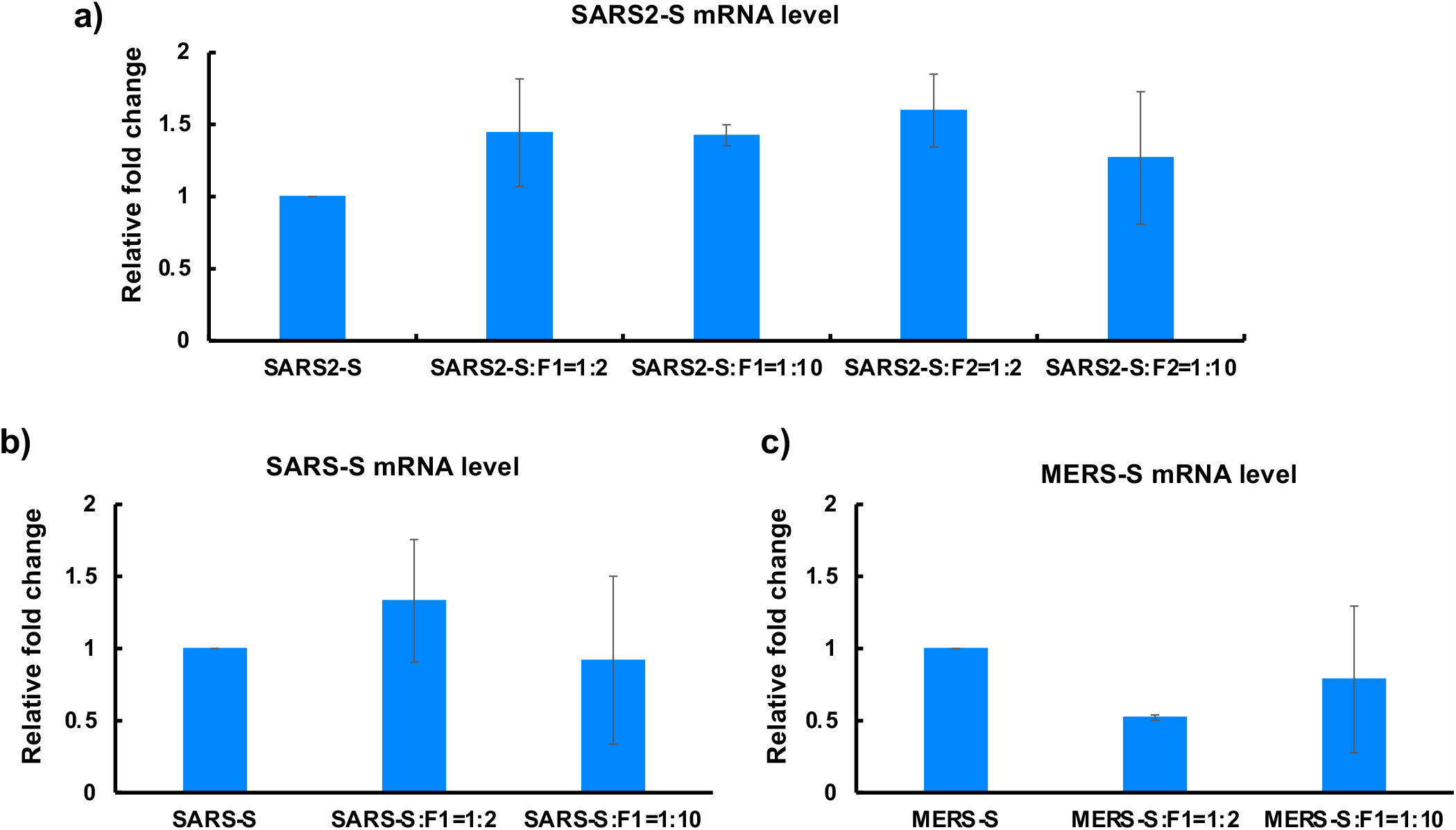
The mRNA levels of coronavirus spike proteins in the absence or presence of polypeptide interference. **a)**. COVID-19 SARS2-S; **b)**. 2002 SARS-S; **c). 2**012 MERS-S.

**Figure S6.**
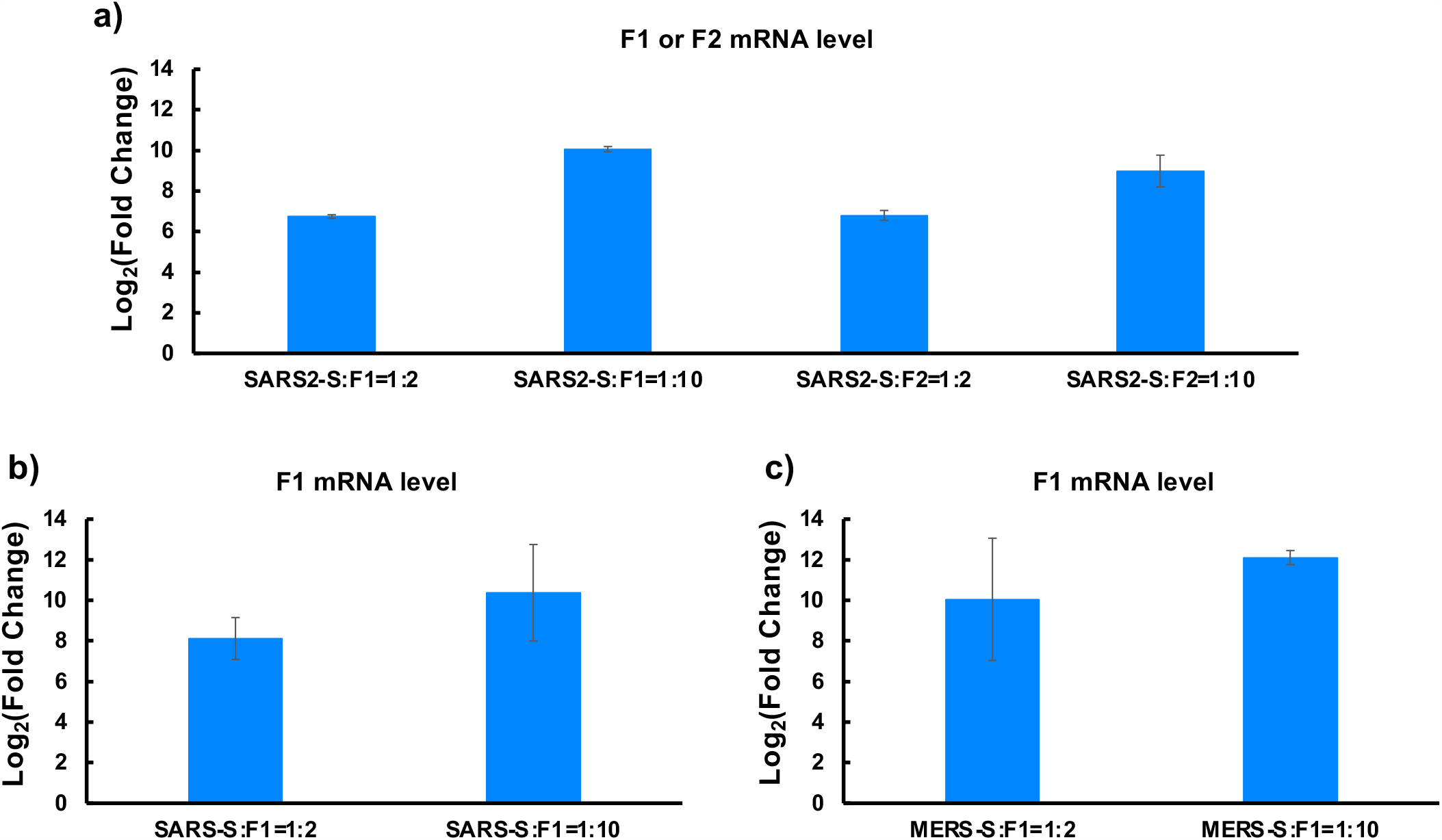
The mRNA levels of F1 and F2 when co-transfected with coronavirus spike protein-coding plasmids. **a)**. with COVID-19 SARS2-S; **b)**. with 2002 SARS-S; **c)**. with 2012 MERS-S.

**Figure S7.**
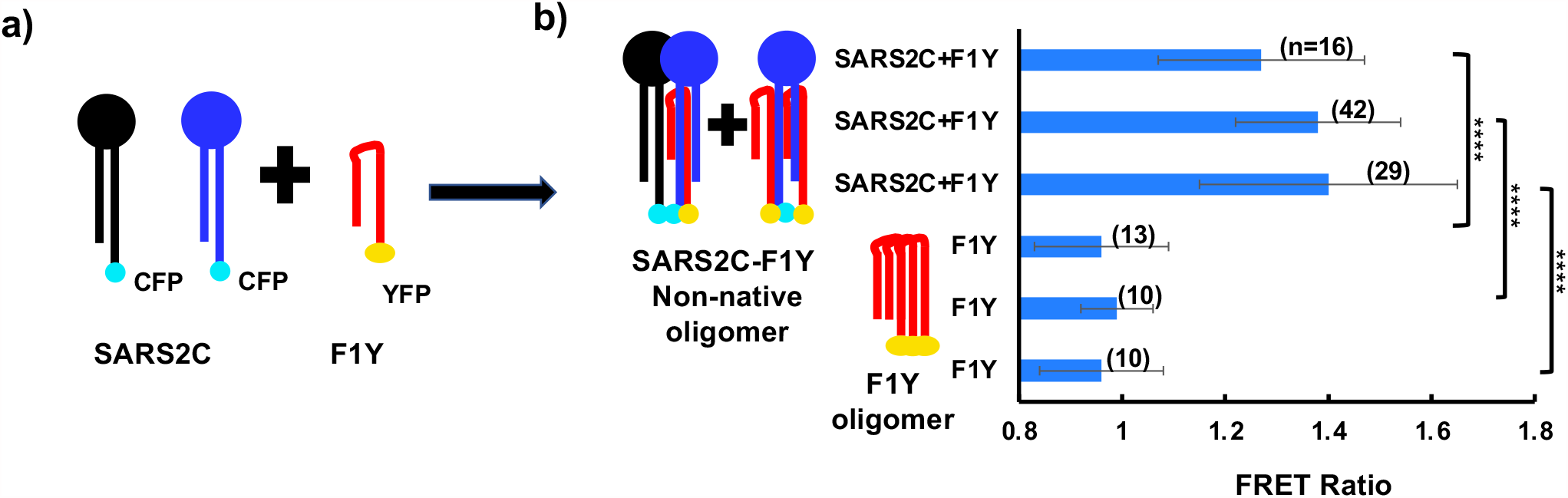
F1 interacted with coronavirus spike proteins at the protein level. **a)**. Diagram for the FRET setup, where CFP and YFP were each tagged to the extreme C-terminus of SARS2-S and F1, resulting in SARS2C and F1Y, respectively. **b)**. FRET ratio of SARS2C with F1Y (at a 1:1 molar ratio for transfection), compared to that of F1Y only. ****, *p*<0.0001 in unpaired *t*-test comparing FRET ratios of SARS2C+F1Y *versus* same-day F1Y only. Numbers in parentheses indicate the number of cells included in each analysis.

## References

1 V’Kovski, P., Kratzel, A., Steiner, S., Stalder, H. & Thiel, V. Coronavirus biology and replication: implications for SARS-CoV-2. Nature reviews. Microbiology 19, 155–170, doi:10.1038/s41579-020-00468-6 (2021).

2 Rambaut, A. et al. Preliminary genomic characterisation of an emergent SARS-CoV-2 lineage in the UK defined by a novel set of spike mutations. (2020).

3 Tegally, H., Wilkinson, E. & al., e. Emergence and rapid spread of a new severe acute respiratory syndrome-related coronavirus 2 (SARS-CoV-2) lineage with multiple spike mutations in South Africa. medRxiv 2020.2012.2021.20248640 (2020).

4 Naveca, F. et al. Phylogenetic relationship of SARS-CoV-2 sequences from Amazonas with emerging Brazilian variants harboring mutations E484K and N501Y in the Spike protein. virological.org (2021).

5 Zhou, D., Dejnirattisai, W., Supasa, P., Liu, C. & al., e. Evidence of escape of SARS-CoV-2 variant B.1.351 from natural and vaccine induced sera. Cell In press (2021).

6 Tada, T. et al. Neutralization of viruses with European, South African, and United States SARS-CoV-2 variant spike proteins by convalescent sera and BNT162b2 mRNA vaccine-elicited antibodies. bioRxiv, doi:10.1101/2021.02.05.430003 (2021).

7 Wang, P. et al. Increased Resistance of SARS-CoV-2 Variants B.1.351 and B.1.1.7 to Antibody Neutralization. bioRxiv, doi:10.1101/2021.01.25.428137 (2021).

8 Zhou, B. et al. SARS-CoV-2 spike D614G change enhances replication and transmission. Nature, doi:10.1038/s41586-021-03361-1 (2021).

9 Cele, S. et al. Escape of SARS-CoV-2 501Y.V2 from neutralization by convalescent plasma. Nature, doi:10.1038/s41586-021-03471-w (2021).

10 Wrapp, D. et al. Cryo-EM structure of the 2019-nCoV spike in the prefusion conformation. Science 367, 1260–1263, doi:10.1126/science.abb2507 (2020).

11 Walls, A. C. et al. Structure, Function, and Antigenicity of the SARS-CoV-2 Spike Glycoprotein. Cell 183, 1735, doi:10.1016/j.cell.2020.11.032 (2020).

12 Shang, J. et al. Structural basis of receptor recognition by SARS-CoV-2. Nature 581, 221–224, doi:10.1038/s41586-020-2179-y (2020).

13 Cai, Y. et al. Distinct conformational states of SARS-CoV-2 spike protein. Science 369, 1586–1592, doi:10.1126/science.abd4251 (2020).

14 Bangaru, S. et al. Structural analysis of full-length SARS-CoV-2 spike protein from an advanced vaccine candidate. Science 370, 1089–1094, doi:10.1126/science.abe1502 (2020).

15 Klein, S. et al. SARS-CoV-2 structure and replication characterized by in situ cryo-electron tomography. Nat Commun 11, 5885, doi:10.1038/s41467-020-19619-7 (2020).

16 Stertz, S. et al. The intracellular sites of early replication and budding of SARS-coronavirus. Virology 361, 304–315, doi:10.1016/j.virol.2006.11.027 (2007).

17 Coutard, B. et al. The spike glycoprotein of the new coronavirus 2019-nCoV contains a furin-like cleavage site absent in CoV of the same clade. Antiviral Res 176, 104742, doi:10.1016/j.antiviral.2020.104742 (2020).

18 Johnson, B. A. et al. Loss of furin cleavage site attenuates SARS-CoV-2 pathogenesis. Nature 591, 293–299, doi:10.1038/s41586-021-03237-4 (2021).

19 Cui, J., Li, F. & Shi, Z. L. Origin and evolution of pathogenic coronaviruses. Nature reviews. Microbiology 17, 181–192, doi:10.1038/s41579-018-0118-9 (2019).

20 de Wit, E., van Doremalen, N., Falzarano, D. & Munster, V. J. SARS and MERS: recent insights into emerging coronaviruses. Nature reviews. Microbiology 14, 523–534, doi:10.1038/nrmicro.2016.81 (2016).

21 Erickson, M. G., Alseikhan, B. A., Peterson, B. Z. & Yue, D. T. Preassociation of calmodulin with voltage-gated Ca(2+) channels revealed by FRET in single living cells. Neuron 31, 973–985, doi:10.1016/s0896-6273(01)00438-x (2001).

22 Hardee, C. L., Arevalo-Soliz, L. M., Hornstein, B. D. & Zechiedrich, L. Advances in Non-Viral DNA Vectors for Gene Therapy. Genes (Basel) 8, doi:10.3390/genes8020065 (2017).

23 Catanese, D. J., Jr., Fogg, J. M., Schrock, D. E., 2nd, Gilbert, B. E. & Zechiedrich, L. Supercoiled Minivector DNA resists shear forces associated with gene therapy delivery. Gene Ther 19, 94–100, doi:10.1038/gt.2011.77 (2012).

24 Kay, M. A., He, C. Y. & Chen, Z. Y. A robust system for production of minicircle DNA vectors. Nat Biotechnol 28, 1287–1289, doi:10.1038/nbt.1708 (2010).

25 Zhao, G. et al. A safe and convenient pseudovirus-based inhibition assay to detect neutralizing antibodies and screen for viral entry inhibitors against the novel human coronavirus MERS-CoV. Virol J 10, 266, doi:10.1186/1743-422X-10-266 (2013).

